# Towards identifying subnetworks from FBF binding landscapes in *Caenorhabditis* spermatogenic or oogenic germlines

**DOI:** 10.1101/422584

**Authors:** Douglas F. Porter, Aman Prasad, Brian H. Carrick, Peggy Kroll-Connor, Marvin Wickens, Judith Kimble

## Abstract

Metazoan PUF (Pumilio and FBF) RNA-binding proteins regulate various biological processes, but a common theme across phylogeny is stem cell regulation. In *Caenorhabditis elegans*, FBF (*fem-3* Binding Factor) maintains germline stem cells regardless of which gamete is made, but FBF also functions in the process of spermatogenesis. We have begun to “disentangle” these biological roles by asking which FBF targets are gamete-independent, as expected for stem cells, and which are gamete-specific. Specifically, we compared FBF iCLIP binding profiles in adults making sperm to those making oocytes. Normally, XX adults make oocytes. To generate XX adults making sperm, we used a *fem-3(gf)* mutant requiring growth at 25°; for comparison, wild-type oogenic hermaphrodites were also raised at 25°. Our FBF iCLIP data revealed FBF binding sites in 1522 RNAs from oogenic adults and 1704 RNAs from spermatogenic adults. More than half of these FBF targets were independent of germline gender. We next clustered RNAs by FBF-RNA complex frequencies and found four distinct blocks. Block I RNAs were enriched in spermatogenic germlines, and included validated target *fog-3*, while Block II and III RNAs were common to both genders, and Block IV RNAs were enriched in oogenic germlines. Block II (510 RNAs) included almost all validated FBF targets and was enriched for cell cycle regulators. Block III (21 RNAs) was enriched for RNA-binding proteins, including previously validated FBF targets *gld-1* and *htp-1*. We suggest that Block I RNAs belong to the FBF network for spermatogenesis, and that Blocks II and III are associated with stem cell functions.

## INTRODUCTION

RNA regulatory networks — defined by genome-wide interactions between RNA-binding proteins and their RNA targets — are central to biological control (Keene 2007; Ascano *et al.* 2013; Ule and Darnell 2006; Ivshina *et al.* 2014). Among RNA-binding proteins analyzed at a genomic level for target RNAs, the PUF RNA-binding proteins (for <u>Pu</u>milio and <u>F</u>BF) have served as paradigms because of exquisite sequence-specificity and high affinity for their binding elements (Wang *et al.* 2001; Wang *et al.* 2002; Wang *et al.* 2009; Qiu *et al.* 2012; Zhu *et al.* 2009). For example, each of five PUF proteins in *Saccharomyces cerevisae* binds a battery of mRNAs, with some redundancy for targets in those networks but with key biological functions associated with each particular PUF (Gerber *et al.* 2004; Porter *et al.* 2015; Wilinski *et al.* 2015). Metazoans also have one or more PUF proteins with multiple biological roles. An ancient and apparently common function of metazoan PUFs is stem cell maintenance (Wickens *et al.* 2002), but PUFs can also regulate sex determination, embryonic polarity, neurogenesis and learning, among their varied biological roles (Lin and Spradling 1997; Spradling *et al.* 2001; Crittenden *et al.* 2002; Wickens *et al.* 2002; Spassov 2004; Salvetti *et al.* 2005; Kaye *et al.* 2009; Vessey *et al.* 2010; Campbell *et al.* 2012; Lander *et al.* 2012; Zhang *et al.* 1997; Zhang *et al.* 2017; Darnell 2013; Follwaczny *et al.* 2017). Moreover, mutations in the human PUM1 gene can lead to both developmental delay and seizures (Gennarino *et al.* 2018). The challenge now is to identify metazoan PUF subnetworks with distinct biological roles and to define those mRNAs whose regulation is critical for stem cells.

The *C. elegans* PUF paralogs, FBF-1 and FBF-2 (collectively known as FBF), are exemplars of metazoan PUF regulation. FBF-1 and FBF-2 are major regulators of germline stem cell maintenance (Crittenden *et al.* 2002), the hermaphrodite sperm-to-oocyte switch (Zhang *et al.* 1997), and the process of spermatogenesis (Luitjens *et al.* 2000). FBF preferentially binds its targets in the 3’UTR in a sequence-specific fashion (Prasad *et al.* 2016). The FBF binding element (FBE) is UGUNNNAU with the optimal FBE being UGUDHHAU, where D is A, U, or G and H is A, U, or C (Bernstein *et al.* 2005; Opperman *et al.* 2005); moreover, cytosine residues located one or two positions upstream of the FBE (−1C or −2C) enhance affinity (Qiu *et al.* 2012). Like most PUF proteins, FBF recruits other proteins to its target mRNAs (Suh *et al.* 2009; Friend *et al.* 2012; Kraemer *et al.* 1999; Luitjens *et al.* 2000; Eckmann *et al.* 2002; Campbell *et al.* 2012; Shin *et al.* 2017) and is best known for decreasing RNA stability or repressing translation (Zhang *et al.* 1997; Crittenden *et al.* 2002; Merritt *et al.* 2008; Zanetti *et al.* 2012; Shin *et al.* 2017); however, FBF can also activate mRNAs (Kaye *et al.* 2009; Suh *et al.* 2009) and has been proposed to mediate the transition from self-renewal to differentiation via a switch from its repressive to its activating mode (Kimble and Crittenden 2007). Consistent with this idea, a regulated transition from PUF-mediated repression to activation was recently found for a yeast PUF (Lee and Tu 2015).

Previous genomic analyses of the network of RNAs associated with FBF-1 and FBF-2 focused on adult oogenic germlines (Kershner and Kimble 2010; Prasad *et al.* 2016). Most relevant to this work were the iCLIP studies showing that FBF-1 and FBF-2 associate with largely the same mRNAs via the same binding sites (Prasad *et al.* 2016). Therefore, FBF-1 and FBF-2 are not only biologically redundant for regulation of stem cells (Crittenden *et al.* 2002), but these nearly identical proteins also control a common “FBF network”.

Here we use iCLIP to compare FBF-bound RNAs in spermatogenic and oogenic germlines with the goal of identifying subnetworks responsible for individual FBF biological functions. Because FBF is essential for regulation of stem cells in both spermatogenic and oogenic germlines (Crittenden *et al.* 2002), we reasoned that identification of gamete-independent FBF target mRNAs might help define the FBF stem cell network. Conversely, spermatogenic-specific FBF target mRNAs might represent the FBF subnetwork responsible for spermatogenesis. We combine experimental and computational approaches to identify likely FBF targets and to propose subnetworks.

## MATERIALS AND METHODS

### Nematode strains used in this study

JK4561: *fem-3(q22 ts,gf) IV*

JK5181: *fbf-1(ok91) qSi232[3xflag::fbf-1] II*

JK5182: *fbf-2(q738) qSi75[3xflag::fbf-2] II*

JK5140: *fbf-1(ok91) qSi232[3xflag::fbf-1] II; fem-3(q22 ts,gf) IV*/ *nT1[qIs51](IV;V)*

JK5545: *fbf-2(q738) qSi75[3xflag::fbf-2] II*; *fem-3(q22 ts,gf) IV*/ *nT1[qIs51](IV;V)*

### Generation and maintenance of strains carrying epitope-tagged FBF-1 and FBF-2 transgenes

Strains JK5181 and JK5182 were generated previously (Prasad *et al.* 2016). Briefly, the *qSi232* (3xFLAG::FBF-1) and *qSi75* (3xFLAG::FBF-2) transgenes were created by the method of *Mos1*-mediated single copy insertion (MosSCI) (Frøkjær-Jensen *et al.* 2008) and placed into strains lacking *fbf-1* or *fbf-2* respectively. Like wild-type, these strains are oogenic as adults; the primary difference is that they carry a FLAG-tagged FBF so that we can do iCLIP. To generate spermatogenic adults, we used genetic crosses to introduce the temperature gain-of-function allele, *fem-3(q22 ts,gf)*, and thereby generated JK5140 and JK5545. This *fem-3* mutant is spermatogenic when grown at 25° from the first larval stage (L1) (Barton *et al.* 1987), and strains generated with tagged FBF were similarly spermatogenic at 25°. This *fem-3* allele is a T-to-C mutation near one of two 3’UTR FBEs (CGCTTCT<u>TGTGTCAT</u> to CGCT*<u>C</u>*CT<u>TGTGTCAT</u>; FBE underlined, mutation underlined and italicised). To compare oogenic and spermatogenic iCLIP datasets, both oogenic and spermatogenic animals were maintained at 15° for propagation and shifted to 25° from the L1 stage for iCLIP.

### iCLIP

iCLIP was carried out with modifications for *C. elegans,* as previously described (Huppertz *et al.* 2014; Prasad *et al.* 2016). Single-end sequencing was performed on an Illumina HiSeq 2000. Data is available in the NCBI GEO database, accession GSE83695.

### Data processing

Fastq files were split by barcode and 3’ linker and reverse transcription primer sequences were clipped. The barcode was then removed from the read sequence and moved to the read name. Reads were mapped to the genome using STAR and CSEQ parameters (Dobin *et al.* 2013; Kassuhn *et al.* 2016), except alignment was local, rather than end-to-end. Reads mapping to multiple places by STAR or mapping with a STAR-reported score below 20 were removed. Duplicates were removed using scripts from Weyn-Vanhentenryck *et al.* (2014) applied to the barcode sequence found in read names. Reads were assigned to RNAs by HTSeq (Anders *et al.* 2015) and initial differential expression analysis was performed by DESeq2 (v. 1.181.). Peaks were called as described previously (Prasad *et al.* 2016), except that two reads-in-peak cutoffs were applied: one cutoff by per-million normalized read number (2-fold or higher), and one by un-normalized read number (5-fold or higher). The exact cutoffs varied between datasets and are included in File S2. We used the same criteria as described previously to determine cutoffs (Prasad *et al.* 2016). Specifically, cutoffs were chosen to retain all validated targets, maximize enrichment of the binding site, and identify as many potential targets as possible. HOMER (Heinz *et al.* 2010) was performed using the highest 500 peaks, with the single parameter “-rna”.

### Generation of “FBF” replicates for 25° datasets

One FBF-1 biological replicate from 25° oogenic worms had fewer reads than the other five for this strain (two for FBF-1 and three for FBF-2), although the other FBF-1 replicates are large enough that the FBF-1 dataset is still larger than the FBF-1 dataset (Figure S1A, File S2). To generate replicates of more comparable size, we combined iCLIP reads for FBF-1 and FBF-2 to generate three more equally sized FBF replicates. We did not face a similar problem with replicates from spermatogenic animals (Figure S1B), but similarly combined these as well.

### Clustering method

We first normalized each of our iCLIP datasets to reads-per-million so that reads-per-RNA represented the frequency of binding at a given RNA. We then subtracted the average of the negative controls from each FBF iCLIP dataset, and finally converted all counts to a log**_2_** scale. We calculated distances between binding frequencies for each FBF-RNA pair by Euclidean distance and clustered those distances by simple hierarchial clustering (Eisen *et al.* 1998). Euclidean distance is a generalization of the notion of distance as the length of a straight line between two points. In our case, the distance between two RNAs *A* and *B* is the distance between the vectors of FBF binding (each FBF iCLIP replicate being one dimension of the vector) at *A* and *B*. We used Euclidean distance between reads-per-million counts, rather than normalizing each RNA to have the same average number of reads, so that we could cluster according to both frequency of binding and the dependency of binding on germline gender. Distance metrics were used to generate clusters using pairwise average-linkage cluster analysis (Sokal and Michener 1958), in which distances between clusters are simply defined as the average of all distances between elements in a cluster *A* with all elements in a cluster *B*.

### DESeq2

A read was assigned to an RNA if and only if it overlapped with an exon of the corresponding gene. DESeq2 (v. 1.18.1) results were generated as described in the DESeq2 documentation, using default parameters of the DESeq function, which set minimum read depths based on maximizing the genes passing a given FDR. We used an FDR of 0.01. DESeq-reported p-values are Benjamini-Hochberg-adjusted. We applied DESeq2 analysis to compare the effect of both gender and of temperature. In either case, we restricted our analysis to those RNAs identified as part of the spermatogenic or oogenic program (Noble *et al.* 2016).

### Worm-human PUF target comparison

We used a compendium of *C. elegans* genes with human orthologs (Shaye and Greenwald 2011) to identify which FBF target RNAs encode proteins with human counterparts. We compared these FBF targets to PUM2 targets identified by PAR-CLIP in human embryonic kidney cells (Hafner *et al.* 2010) and to PUM1 and PUM2 targets identified by iCLIP during mouse neurogenesis (Zhang *et al.* 2017). An FBF target was defined as shared with PUM if (1) any mammalian ortholog was targeted by PUM, (2) there were no more than ten mammalian orthologs (such limits have been used previously for cross-phyla comparison (Hogan *et al.* 2015)), and (3) there were no more than ten *C. elegans* genes orthologous to the same mammalian ortholog. We treated orthology as a transitive property: if nematode genes “A” and “B” are listed as orthologs in Shaye and Greenwald (2011) to mammalian genes that overlap by at least one gene, then “A” and “B” were treated as if they were a single gene for calculating overlap. The same method of combining orthologs was applied to the mammalian gene set (if two mammalian genes overlap in worm orthologs, they were combined).

### Statistical analysis

All statistical methods for determining FBF-RNA interactions were as described in Prasad *et al.* (2016). Briefly, reads in the 500-bp region around a peak were placed in 50-bp bins for both FBF iCLIP and no-antibody iCLIP control data. The negative control was modelled as a Gaussian to calculate a p-value as the chance of observing a peak at the given height from the negative control data. All p-values were then Benjamini-Hochberg corrected and an FDR cutoff of 1% applied, before applying the two ratio cutoffs described above. Statistics used for DESeq2 fold-change estimates and target comparison are described above.

### Data Availability

Strains are available upon request. Scripts used to analyze the data were uploaded to github.com/dfporter/FBF_gendered_gl. Sequencing data is available in the NCBI GEO database, accession GSE83695. To replicate the combined 25° FBF datasets from individual FBF-1 and FBF-2 replicates, first obtain the individual replicates from GSE83695, and concatenate 25° oogenic FBF-1/FBF-2 replicates in the order 1/3, 2/2, and 3/1; then concatenate 25° spermatogenic FBF-1/FBF-2 replicates in the order 2/1, 1/2, and 3/3. The 20° FBF iCLIP data from Prasad *et al.* (2016) is available at GSE76136. File S1 contains FBF iCLIP peaks. File S2 contains metrics such as complexity for FBF iCLIP peaks. File S3 contains GO terms for FBF targets. File S4 describes RNAs significantly differing between spermatogenic and oogenic in FBF iCLIP. File S5 contains the dataset displayed in Figure 3A, namely FBF binding per gene for 2,111 FBF target RNAs. File S6 contains the blocks defined in Figure 3. Finally, File S7 contains FBF targets overlapping with the human PUF protein PUM2.

## RESULTS AND DISCUSSION

### Generation of FBF iCLIP datasets from spermatogenic and oogenic germlines

We generated FBF-1 and FBF-2 iCLIP datasets from animals with somatic tissues of the same gender but germline tissue of opposite gender (Figure 1, A and B). All animals were chromosomally XX and had hermaphroditic somatic tissues, including the somatic gonad; they also had comparable numbers of germline stem cells but those stem cells generated either only oocytes or only sperm, depending on the strain. For each FBF, we used an N-terminal 3XFLAG-tagged single copy transgene in a strain lacking the endogenous gene (e.g. FLAG::FBF-1 in an *fbf-1* null mutant). As reported before (Prasad *et al.* 2016), *fbf(0)* FLAG::FBF XX animals are essentially wild-type. Moreover, each tagged FBF rescues *fbf-1 fbf-2* double mutants from 100% sterility due to lack of GSCs to 100% fertility due to rescue of the GSC defect (Prasad *et al.* 2016). These tagged FBFs should therefore interact in an essentially normal fashion with their target RNAs.

**Figure 1.**
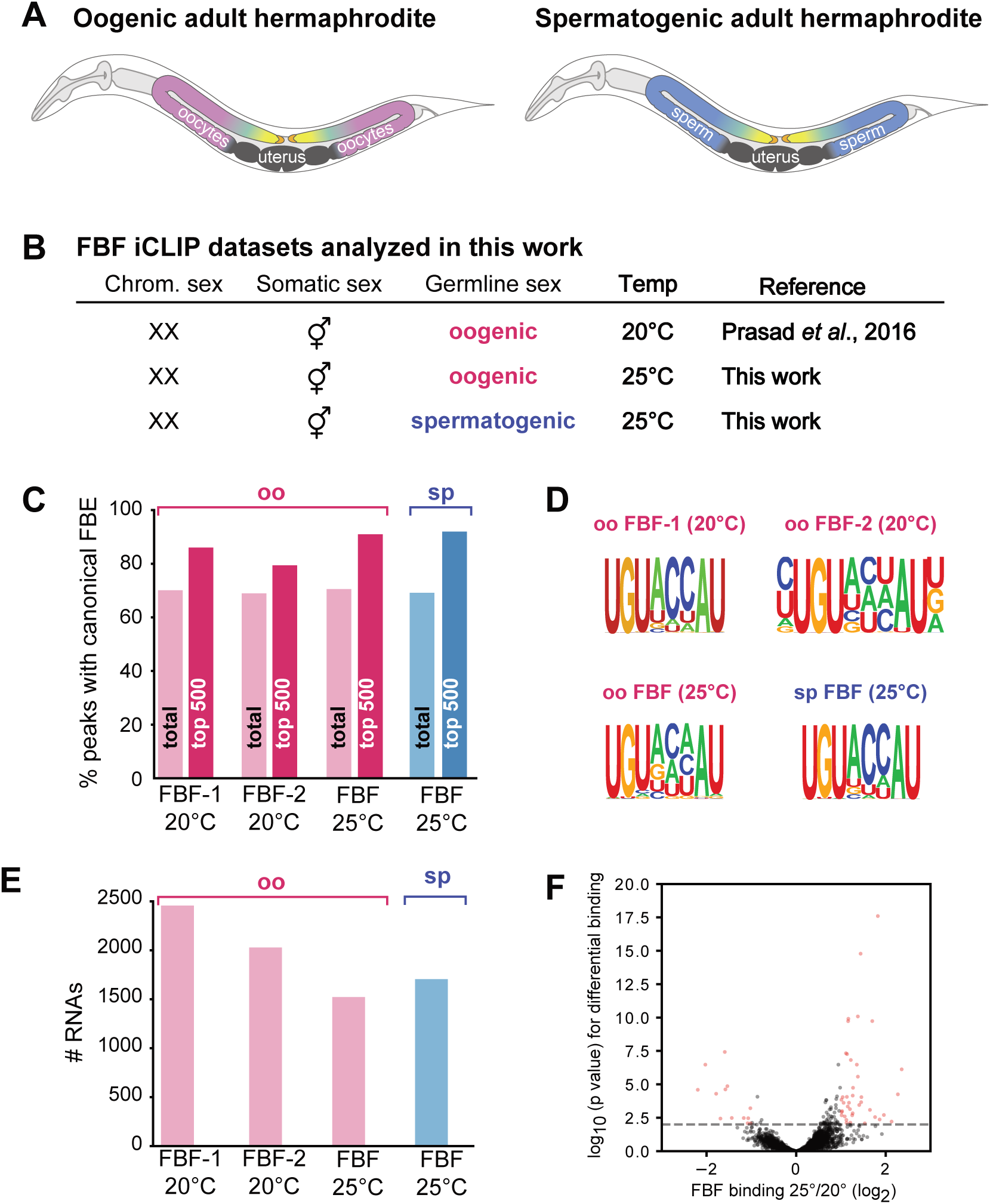
(A) Diagrams of adult XX hermaphrodites making only oocytes (left) or only sperm (right), but with hermaphroditic somatic tissues. Somatic tissues, grey; oogenic germline, rose; spermatogenic germline, blue. Germline stem cells (yellow) exist in both oogenic and spermatogenic germlines, and are maintained by signaling from their niche (orange). (B) FBF iCLIP datasets analyzed in this work. (C) Percentage of peaks containing a canonical FBE (UGUNNNAU) in FBF iCLIP datasets. oo, iCLIP from oogenic worms; sp, iCLIP from spermatogenic worms; 20° or 25°, temperature at which worms were raised. For each dataset, we scored all peaks (bars marked “total”) as well as the top 500 peaks (bars marked “top 500”). (D) For each dataset, the canonical FBE was the most significant motif in the top 500 FBF peaks, according to HOMER. (E) Number of distinct FBF target RNAs identified for indicated datasets. (F) Few RNAs are differentially bound by FBF between oogenic worms raised at 25° and 20°, as judged by DESeq2 analysis of reads-per-gene for 5,768 genes with an average of least 20 reads in oogenic FBF iCLIP and which are expressed in the germline (Noble *et al.* 2016). The x-axis denotes the fold change of FBF binding (reads-per-gene) in 25° worms over 20° worms, while the y-axis denotes the statistical significance of differential binding. The dashed line is indicates a P value of 0.01. Red dots are the 1% (54) of genes with >2 fold change and P value < 0.01.

XX adults with a spermatogenic germline were obtained using a temperature sensitive gain-of-function *(gf) fem-3* mutant (Barton *et al.* 1987). We crossed transgenes encoding 3XFLAG-tagged FBF-1 or FBF-2 into the *fem-3(gf)* mutant strain, and again removed the corresponding endogenous *fbf* gene in each strain. As expected, the final strains, *fbf-1(0) FLAG::FBF-1; fem-3 (gf)* and *fbf-2(0) FLAG::FBF-2; fem-3 (gf),* were self-fertile at permissive temperature (15°), but fully spermatogenic at restrictive temperature (25°). Because the previously reported oogenic FBF iCLIP was done with animals raised at 20° (Prasad *et al.* 2016), we repeated it here with animals grown at 25°. Thus, we performed FBF iCLIP from adults that were either oogenic or spermatogenic, both raised at 25°. For each strain (each FBF, each germline gender), we processed three biological replicates. In parallel, we produced three negative control replicates for each germline gender by omitting the FLAG antibody from the beads during immunopurification.

### Targets, networks and subnetworks: definitions

Throughout this work, we define the term “target RNAs” empirically as RNAs that interact with FBF after cross-linking in living animals, followed by immunoprecipitation from lysate and deep sequencing (CLIP). We define “network” to encompass all RNA targets observed by CLIP, and “sub-network” as a subset of that broader network. We refer to RNAs whose expression is regulated by FBF as “validated targets”. Such validation relies on genetic, biochemical and cellular analyses that have been done by ourselves and others in previous studies. We note that virtually all validated FBF targets are among the targets identified in this work by FBF CLIP (see below).

### Peak calling and generation of quality datasets for comparison

This work takes advantage of three sets of FBF iCLIP data (Figure 1B). To analyze these datasets, we modified our earlier peak calling pipeline (Prasad *et al.* 2016) to include a step that collapses duplicate reads while accounting for sequencing errors (Weyn-Vanhentenryck *et al.* 2014) (see Materials and Methods). This modified pipeline generated lists of FBF-1 and FBF-2 target RNAs in oogenic animals raised at 25° and spermatogenic animals raised at 25°, as well as revised lists of FBF-1 and FBF-2 targets in oogenic animals raised at 20°. File S1 lists iCLIP peaks obtained for each condition, and File S2 presents metrics of dataset size and quality.

The primary motivation for this work was comparison of FBF targets in spermatogenic and oogenic germlines, with the goal of identifying gamete-independent and gamete-specific targets that might inform about FBF subnetworks. Such comparisons are best done with datasets of comparable size. For iCLIP data of animals raised at 25°, we initially called peaks for FBF-1 and FBF-2 separately (File S1, Figure S1A-C), but one 25° FBF-1 replicate from oogenic germlines had a low number of unique and uniquely mapping reads (13,486, File S2). Because of the similarity of FBF-1 and FBF-2 binding (Prasad *et al.* 2016; this work) and the increased sensitivity of using larger datasets, we combined the FBF-1 and FBF-2 25° iCLIP datasets to generate “FBF” datasets for each gender (see Materials and Methods). Although the differences between FBF-1 and FBF-2 merit future investigation, combining datasets allowed us to more easily compare FBF binding at 25° between genders. These FBF target lists comprised 1,522 RNAs for oogenic animals, and 1,704 RNAs for spermatogenic animals (Figure 2A, File S1).

**Figure 2.**
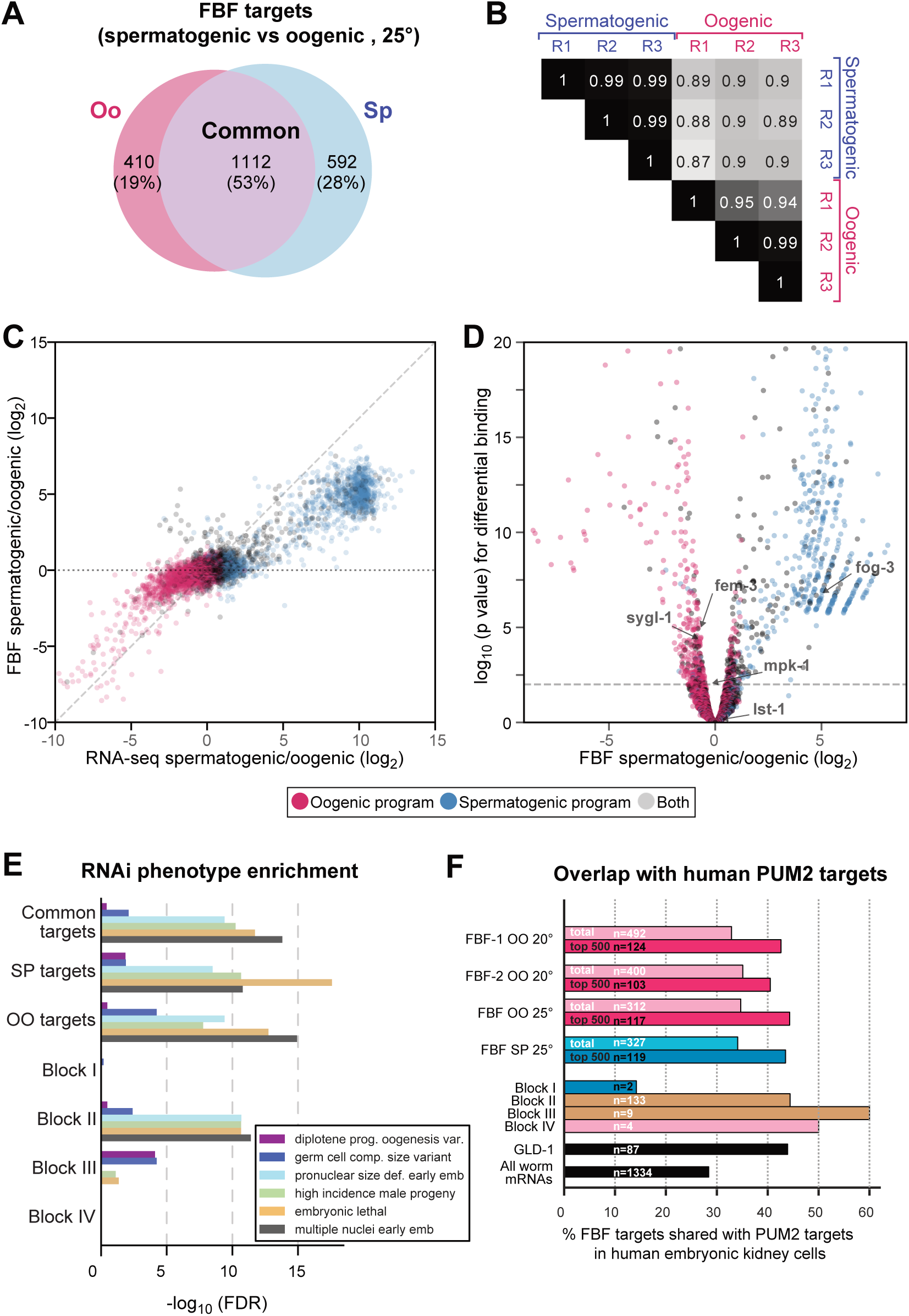
Comparisons of FBF target RNAs in spermatogenic and oogenic germlines. (A) Comparison of FBF targets in spermatogenic (blue) and oogenic animals (pink), both raised at 25°. The 1,112 common, 592 spermatogenic and 410 oogenic represent a first glimpse of potential FBF subnetworks. (B) Spearman correlations between iCLIP replicates (R) reinforce the conclusion that FBF has distinct binding landscapes in spermatogenic and oogenic germlines. Numbers represent rho values for the Spearman correlation between replicates. The oogenic FBF replicate R1 was of lower complexity than the others, which likely explains its lower correlations. (C) Spermatogenic/oogenic ratios of FBF binding (y-axis) to spermatogenic/oogenic ratios of RNA abundance (x-axis). Each dot represents an RNA: pink dots, oocyte-specific RNAs, blue dots, spermatogenic specific RNAs, and grey dots, RNA present in both genders, with germline gender-specificity assigned according to (Noble *et al.* 2016). Dots represent all RNAs with at least one read in any of our datasets, and present in the spermatogenic or oogenic RNA program (Noble *et al.* 2016). Diagonal dashed line, a perfect correlation with slope 1; horizontal dotted line, no correlation. (D) Differences in FBF binding between spermatogenic and oogenic germlines. Color coding of pink, blue and grey is same as in panel (C). 12% of RNAs change binding-frequency significantly (p<0.05, 2-fold) between genders, most of which are enriched in spermatogenic germlines (right arm of volcano plot has more dots than left arm). For ease of viewing, this plot cuts out the few RNAs with extreme p-values, which extend to 10^-89^ for spermatogenesis-enriched FBF targets and to 10^-94^ for oogenesis-enriched FBF targets. (E) Enrichment of RNAi phenotypes in indicated groups of RNAs, as measured by significance (Fisher’s exact test). From the WormBase database, the RNAi phenotype labels are described as follows: *Diplotene progression during oogenesis variant* = developing oocytes are defective during the diplotene stage of meiosis; *Germ cell compartment size variant* = change in germ cell compartment size; *Pronuclear size defective early emb* = size change in pronucleus within gametes or early zygote; *High incidence male progeny* = Higher frequency of male progeny than wild-type; *Embryonic lethal* = progeny die as embryos; *Multiple nuclei early emb* = inviable embryos with more than one nucleus per cell. (F) Overlaps of FBF iCLIP targets with human PUM2 PAR-CLIP targets from human embryonic kidney cells (Hafner *et al.* 2010). The number of ortholog groups comprising the overlap is indicated as “n=”. See text and methods for further explanation. For comparison, the overlap with targets of the germline cell fate regulator GLD-1 (Jungkamp *et al.* 2011) are also given.

**Figure 3.**
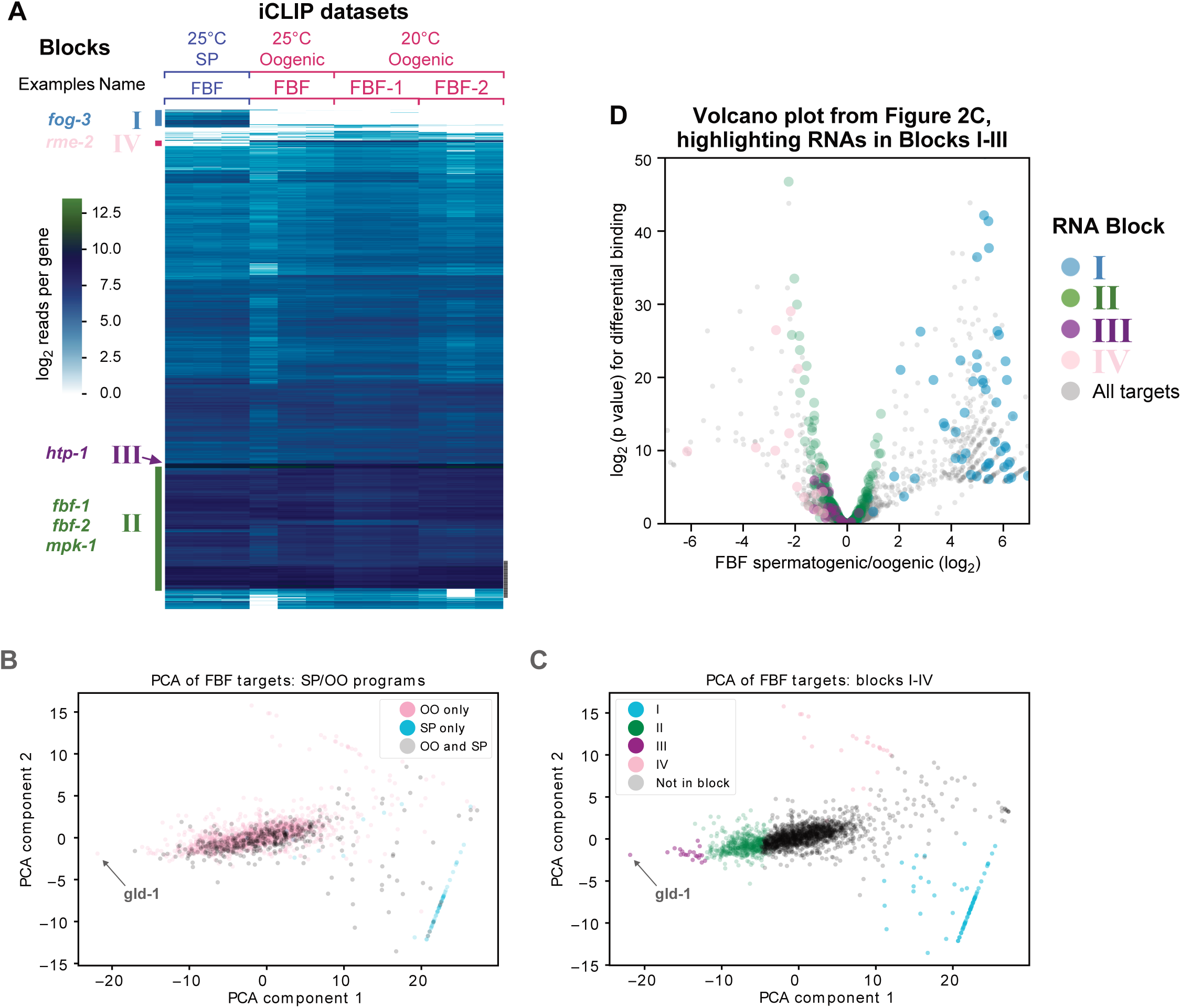
Clustering FBF-RNA complex frequencies reveals four RNA blocks. (A) Columns represent FBF iCLIP samples, as indicated at *top*. Rows represent the 2,114 total RNAs with a significant peak in either of the combined 25° FBF iCLIP datasets. RNAs (rows) were clustered by Euclidean distance and simple hierarchical clustering. Colors represent the log_2_ reads per gene in the given sample (per million reads), after subtracting the negative control. RNA blocks are indicated with Roman numerals. Block I RNAs are enriched in spermatogenic datasets. Block II RNAs are frequently bound and include most established, positive control targets. Block III RNAs are bound at particularly high frequency across all samples. Block IV RNAs are enriched in oogenic datasets. Key examples for each block are noted on left at their approximate location in the y-axis of the heatmap. All positive control RNAs fell into a block except *syp-3*. (B) Principle component analysis of the same FBF-RNA complex frequencies as panel (A) shows RNAs separated by SP/OO ratio (y-axis) and overall FBF-RNA binding frequency (x-axis). RNAs are colored by whether they are only in the oogenic program (pink), only in the spermatogenic program (blue), or in both (grey). A very frequent FBF target RNA across all conditions, *gld-1*, is labeled. (C) The blocks identified in panel (A) are again clustered by PCA, as in (B), but here RNAs are colored by block rather than program. (D) Volcano plot of RNAs from Figure 2D, but color coded by block.

The quality of the datasets analyzed in this work was high by two key criteria. First, the majority of peaks in each dataset contained the canonical FBE (UGUNNNAU), a percentage that rose to roughly 90% for the top 500 peaks (Figure 1C). An “optimal” form of the FBE is an upstream “C” followed by UGURCCAUR, where “R” represents a purine (Prasad *et al.* 2016). Indeed, HOMER identified the FBE as the most enriched motif in the top 500 peaks from all datasets, and a preference for RCC was observed in the degenerate three internal nucleotides, matching the optimal motif (Figure 1D). Second, these target lists include all expected experimentally validated FBF targets. FBF targets in oogenic germlines included 13/15 validated FBF targets (*fbf-1, fbf-2, fem-3, fog-1, gld-1, gld-3, him-3, htp-1, htp-2, syp-2, syp-3, lip-1,* and *mpk-1*), but were missing the two not expected: *fog-3* is sperm-specific and therefore not expressed in oogenic germlines (Chen and Ellis 2000), and *egl-4* has only been established as an FBF target in neurons (Kaye *et al.* 2009) and was not detected in previous genomic analyses of FBF targets (Kershner and Kimble 2010; Prasad *et al.* 2016). Similarly, FBF targets in spermatogenic germlines included 14/15 validated targets: all those in oogenic germlines plus *fog-3*. Finally, both size and complexity of the datasets (File S2) were similar to those for CLIP studies of other PUFs (Hafner *et al.* 2010; Freeberg *et al.* 2013; Porter *et al.* 2015; Wilinski *et al.* 2015) and consistent with our previous report on FBF targets in oogenic germlines (Prasad *et al.* 2016). Thus, all target lists include well over a thousand RNAs (Figure 1E).

Our modified peak calling method revises FBF-1 and FBF-2 target lists in oogenic germlines at 20°, but all major conclusions of our previous study (Prasad *et al.* 2016) were confirmed and revised lists were similar in content. An additional, spermatogenic germline-specific lincRNA *linc-36* was identified for the first time in this analysis along with three previously reported lincRNAs (*linc-7, linc-4*, and *linc-29*). As in our initial report, the cell cycle is the most significantly enriched GO term associated with FBF targets in all of our datasets (File S3). The revised 20° lists contain, respectively, 69% and 84% of FBF-1 and FBF-2 targets reported previously, and the overlap between the FBF-1 and FBF-2 lists remained similar (68-83% of each paralog’s target list overlapped, File S2, Figure S1C). Peak heights for FBF-1 and FBF-2 were highly correlated (Pearson R 0.86), similar to that found previously (Pearson R 0.82) (Prasad *et al.* 2016), confirming the considerable molecular redundancy of these two nearly identical paralogs. Thus, FBF-1 and FBF-2 bind to largely the same target RNAs and largely to the same sites within those RNAs, as concluded previously.

As might be expected, temperature affected the FBF binding landscape but many metrics were comparable: (1) reads-per-gene counts correlated well between temperatures (Figure 1F, Figure S2, average Spearman rho 0.94 between 25° and 20° replicates), (2) a variety of additional metrics were similar (File S2), and (3) targets overlapped heavily (Figure S1D). Figure 1F shows the similarity of reads-per-gene counts for FBF binding at 25° *vs* 20° by DESeq2 analysis: the 1% of RNAs that are significant at a P<0.01 and fold change of >2 are indicated in red. Our peak caller detected peaks in more RNAs in the 20° datasets than in the 25° datasets (Figure 1E), because the 20° datasets have more reads (File S2) and our peak caller has greater sensitivity to detect peaks at higher read depths, despite the distribution of reads-per-gene being similar (Figure 1F).

### Germline gender has a strong influence on the FBF binding landscape

We first compared target RNA identities between iCLIP of spermatogenic and oogenic animals, both grown at 25°. Over half of the FBF target RNAs were shared, but significant fractions were also found only in one germline gender or the other (Figure 2A). Differences due to germline gender were thus greater than differences due to temperature (Figure S2, and Figure 2D compared with Figure 1F). Among the 2114 total FBF targets, 2069 were mRNAs and 45 were non-coding RNAs. For mRNA targets, 1092 were common to both genders (53%), 582 were spermatogenic-specific (28%), and 395 were oogenic-specific (19%); for non-coding RNA targets, 20 were common (44%), 10 were spermatogenic-specific (22%), and 15 were oogenic-specific (33%).

We next gauged differences between spermatogenic and oogenic FBF RNA-binding profiles quantitatively. If each iCLIP sequencing read were derived from a single FBF-RNA interaction *in vivo*, then the number of iCLIP reads mapping to a given RNA, as a fraction of all reads, would serve as an estimate of the frequency of FBF-RNA binding at that RNA (Porter *et al.* 2015). Based on this reasoning, we assessed FBF-RNA binding frequency at each target as the number of FBF iCLIP reads (per million) at a given RNA. We then used Spearman’s rank-order correlation coefficients to compare FBF-RNA binding frequencies across all targets (Figure 2B; Figure S2). Comparisons of the 25° datasets revealed that FBF-RNA binding frequencies correlated well among spermatogenic replicates (Figure 2B, mean correlation 0.99) and among oogenic replicates (Figure 2B, mean 0.96), but more poorly between spermatogenic and oogenic replicates (Figure 2B, mean 0.89, two-tailed p-value 10^-7^ indicating significant difference in correlation between genders compared to within genders by t-test). We broadened this analysis to include FBF binding in oogenic germlines at 20° with similar results (Figure S2). We conclude that FBF binding frequencies correlate well for replicates of the same germline gender but are distinct in spermatogenic and oogenic germlines.

One possibility is that gender-specific differences in FBF binding were simply a reflection of underlying RNA abundances. To investigate this possibility, we assessed the correlations between spermatogenic/oogenic ratios in FBF binding and spermatogenic/oogenic ratios in RNA abundance (Figure 2C). RNA abundances were obtained using RNA-seq data from dissected oogenic and spermatogenic gonads (Ortiz *et al.* 2014). The dashed diagonal line in Figure 2C (slope=1) represents the case in which a given fold-difference in a transcript’s abundance between germlines resulted in the same fold-difference in FBF binding frequency with that RNA. The dotted horizontal line (slope=0) represents the case in which FBF binding frequency had no dependence on germline gender or changes in transcript abundance. The data lies between these extremes. FBF binding frequencies changed between germline genders, and mostly in the same direction as RNA abundance changes. However, changes in FBF binding frequencies were not well correlated with changes in RNA abundance (Pearson R 0.64, Spearman 0.69). In other words, FBF-RNA binding frequencies differ markedly between genders, and do not simply reflect differences in RNA abundance. We conclude that this comparative analysis identifies gamete-specific and gender-neutral FBF targets that begin to outline potential subnetworks.

### Gamete-specific FBF-RNA complex frequencies reflect gamete-specific programs

We asked how the potential FBF subnetworks relate to gamete RNA programs, defined by RNA-Seq (Noble *et al.* 2016). Briefly, each program includes gamete-specific RNAs plus gamete-independent RNAs; for example, the full spermatogenic program includes RNAs expressed only in spermatogenic germlines plus those expressed in germlines making either gamete. We note the spermatogenic program was obtained from worms with a *fem-3* gain-of-function allele (to produce adult spermatogenic animals), as in this work. We asked how FBF-RNA complex frequencies (reads-per-RNA) compare with these spermatogenic and oogenic RNA programs. Out of the total of 12,839 RNAs expressed in the germline (Noble *et al.* 2016), 6,873 possessed an average of at least 20 reads per RNA in FBF iCLIP, and 768 (12%) were bound differentially between the two genders by at least two-fold (p<0.01 by DESeq2 (Love *et al.* 2014); Figure 2D; File S4). Figure 2D depicts the agreement between differentially bound FBF-RNA complex frequencies and gamete programs: RNAs in the spermatogenic program (blue) separate from RNAs in the oogenic program (red) when plotted by the ratio of their differential FBF-RNA complex frequencies. Our results are therefore consistent with previous assignment of RNAs to gamete programs. Interestingly, 557 out of 768 differentially bound RNAs are enriched in spermatogenic germlines, indicating that FBF has a more complex interaction network in spermatogenic germlines than oogenic germlines, as it gains more new RNA targets.

### Search for distinct biological roles associated with FBF subnetworks

We next asked if gamete-neutral, spermatogenic-specific and oogenic-specific targets were enriched for either distinct GO terms or germline phenotypes. No striking difference was found (File S3; Figure 2E). Regardless of germline gender, each group of FBF targets was enriched for genes with similar GO terms and phenotypes. Thus, these groups could not be linked to distinct biological roles.

### Conservation of PUF targets

PUF proteins from diverse branches of Eukarya can perform similar biological functions, including stem cell maintenance (Wickens *et al.* 2002). Moreover, previous studies revealed that they share some of the same target mRNAs, including those regulating the cell cycle and programmed cell death (Kershner and Kimble 2010; Prasad *et al.* 2016; Lee *et al.* 2007). To complement those studies with the expanded and refined FBF target datasets reported in this work, we compared them to the PUM2 PAR-CLIP dataset, obtained from human embryonic kidney cells (HEK293) (Hafner *et al.* 2010) and the PUM1 and PUM2 iCLIP data sets, obtained from neonatal mouse brains (Zhang *et al.* 2017). We collapsed orthologous genes to ortholog groups, and discarded ortholog groups that did not exist in both humans and worms (see methods). Comparison with the PUM2 dataset from HEK293 cells showed that 28% of all ortholog groups were shared with PUM2, while 33-35% of ortholog groups targeted by FBF were shared with PUM2 (Figure 2F, File S7). More striking, among the FBF targets in the top 500 peaks, 40-44% were shared (Figure 2F, File S7). FBF targets in spermatogenic and oogenic germlines had similar overlap. By contrast, comparison of FBF targets with PUM1 and PUM2 targets in mouse neonatal brain were less striking, with an overlap of 15-17% overlap with FBF targets, *vs* 14% of all ortholog groups (Figure S3, File S7). We also compared targets of the *C. elegans* RNA-binding protein GLD-1, which controls the differentiation of germline stem cells, with PUM targets (Jungkamp *et al.* 2011). The GLD-1 target dataset was smaller than the FBF target dataset, but had a similar overlap with human targets (Figure 2F), consistent with the overall number of shared targets reflecting similar molecular and biological functions. For both FBF and GLD-1, target RNAs are more abundant than the average RNA (Figure S4), and this likely also contributes to a higher target overlap with PUM2 than with randomly selected worm genes. This delineation of shared targets provides a resource for further studies.

### Clustering FBF-RNA complex frequencies reveals cores of FBF subnetworks

A common method in systems biology is to cluster the expression of genes across conditions to reveal functionally related groups (Eisen *et al.* 1998). Using that logic, we hypothesized that clustering of the FBF-RNA complex frequencies might also reveal functionally related groups. We began with our list of 2,114 FBF target RNAs and clustered their FBF binding frequencies (Figure 3A; File S5) (see Methods). The actual number of RNAs in Figure 3A and File S5 is 2,111, because our peak caller assigns peaks to the ncRNA if a peak overlaps both mRNA and ncRNA, while such reads were discarded as ambigious when counting reads-per-gene. As a result, three ncRNAs that were assigned peaks were dropped in this analysis for having no reads-per-gene, resulting in 2,111 RNAs. Clustering revealed four blocks of interest, numbered in order of increasing spermatogenic to oogenic binding ratio (Figure 3A, File S6). FBF binds Block I RNAs at high frequency in spermatogenic, but not oogenic animals (Figure 3A). By contrast, FBF binds Block II and Block III RNAs at high frequency in both spermatogenic and oogenic animals and hence are gamete-neutral. Finally, FBF binds a small cluster of RNAs in oogenic but not spermatogenic animals (Block IV), and we note this group is smaller than the reciprocal spermatogenic Block I.

We next compared our results from the heatmap to principle component analysis (PCA, Figure 3B-C). The first component (x-axis) roughly corresponds to an average binding frequency across all datasets, and the second component (y-axis) roughly corresponds to the ratio of spermatogenic *vs* oogenic binding. As a result, dots at the top of the graph are in the oogenic program and dots at the bottom are mostly in the spermatogenic program (Figure 3B). The same clusters observed by clustering the heatmap (Figure 3A) were visible in the PCA plot (Figure 3C), supporting the validity of our groupings. We note that the outlier *gld-1*, which is an extremely frequent FBF target, appears as an extreme example of a Block III RNA in the PCA plot (Fig. 3C), so we added it to Block III.

Figure 3D illustrates these clustered blocks of RNAs together with our earlier DESeq2 analysis of FBF-binding. As expected, Block I and Block IV RNAs were differentially bound in spermatogenic and oogenic animals, respectively (blue and pink dots, Figure 3B), while Block II (green dots, Figure 3B) and Block III (purple dots, Figure 3D) RNAs were either gamete-nonspecific or enriched in oogenic germlines. A major difference between Block II and Block III RNAs was number of reads mapping to the average RNA, which was much greater for Block III than for Block II (Figure 4A). We conclude that distinct groups of RNAs emerge by this clustering method. Below we examine each block in turn.

**Figure 4.**
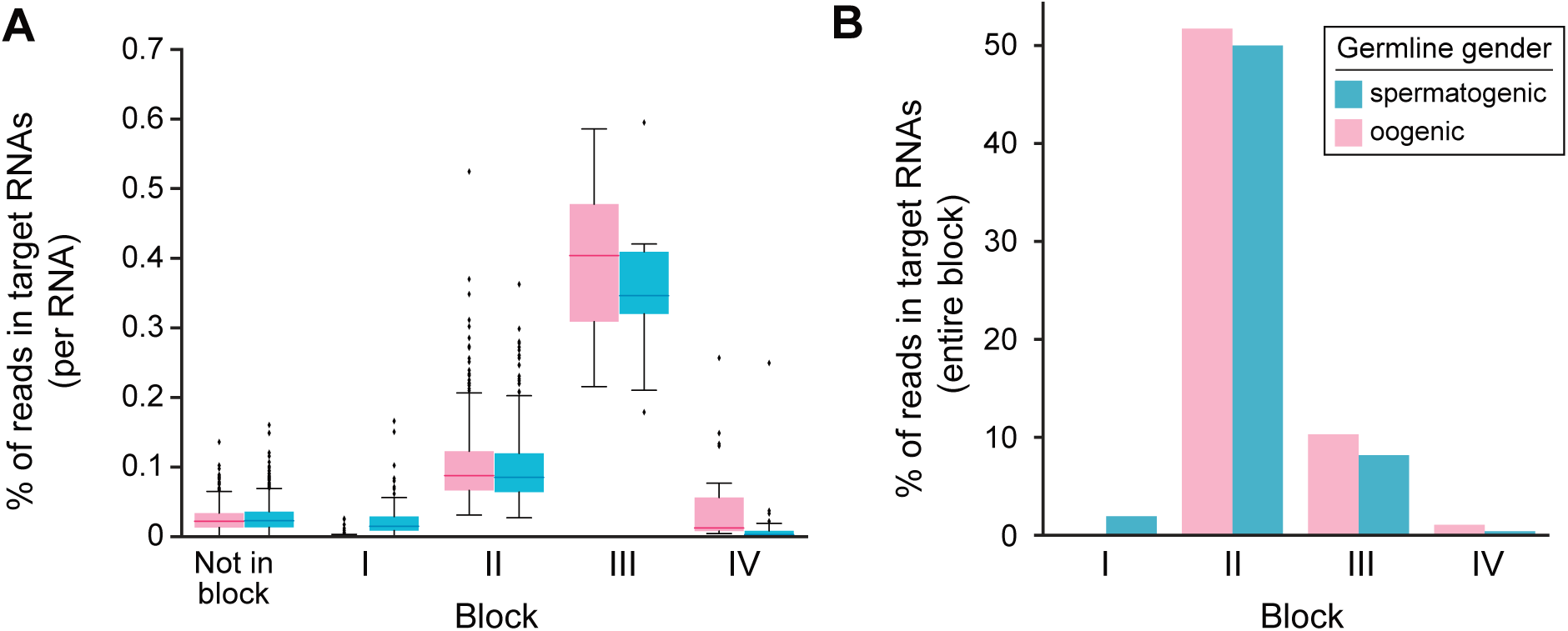
Block RNAs analyzed by germline gender. (A) Percentage of reads mapping to block RNAs. Blue boxplots, reads in spermatogenic animals; pink boxplots, reads in oogenic animals. (B) Percentage of reads in target RNAs from iCLIP in either gender mapping to the indicated block. Thus, roughly 50% of reads in all target RNAs belong to Block II RNAs. The 21 Block III RNAs account for roughly 10% of FBF interactions with target RNAs. Blue bars indicate reads from FBF iCLIP in spermatogenic animals, and pink bars indicate reads from FBF iCLIP in oogenic animals.

### Block I RNAs (File S6)

Block I contains 75 RNAs that are enriched in FBF iCLIP from spermatogenic but not oogenic germlines (Figure 3A). Block I RNAs therefore likely belong to a spermatogenesis FBF subnetwork. Consistent with that idea, Block I RNAs include the key sperm fate regulator *fog-3* (Ellis and Kimble 1995), and 70/75 Block I RNAs belong to the spermatogenic program identified by RNA-seq (Noble *et al.* 2016). However, most Block I RNAs encode proteins whose functions have not yet been characterized and no GO terms were enriched (P value<0.01). To pare down Block I RNAs, we applied two criteria: the highest peak is at least modestly high (25 reads/million) and contains an FBE. This allowed identification of 29 RNAs (Table 1) that encode a diverse array of proteins, some with conserved domains, including a phosphatase and five kinases. This is consistent with FBF serving as a “regulator of regulators” (Kershner and Kimble 2010). We note Block I also contains a novel lncRNA FBF target, *linc-36*. We conclude that Block I RNAs belong to the FBF spermatogenesis subnetwork and that 29 RNAs within Block I are likely to be major FBF targets in that subnetwork.

**Table 1.**
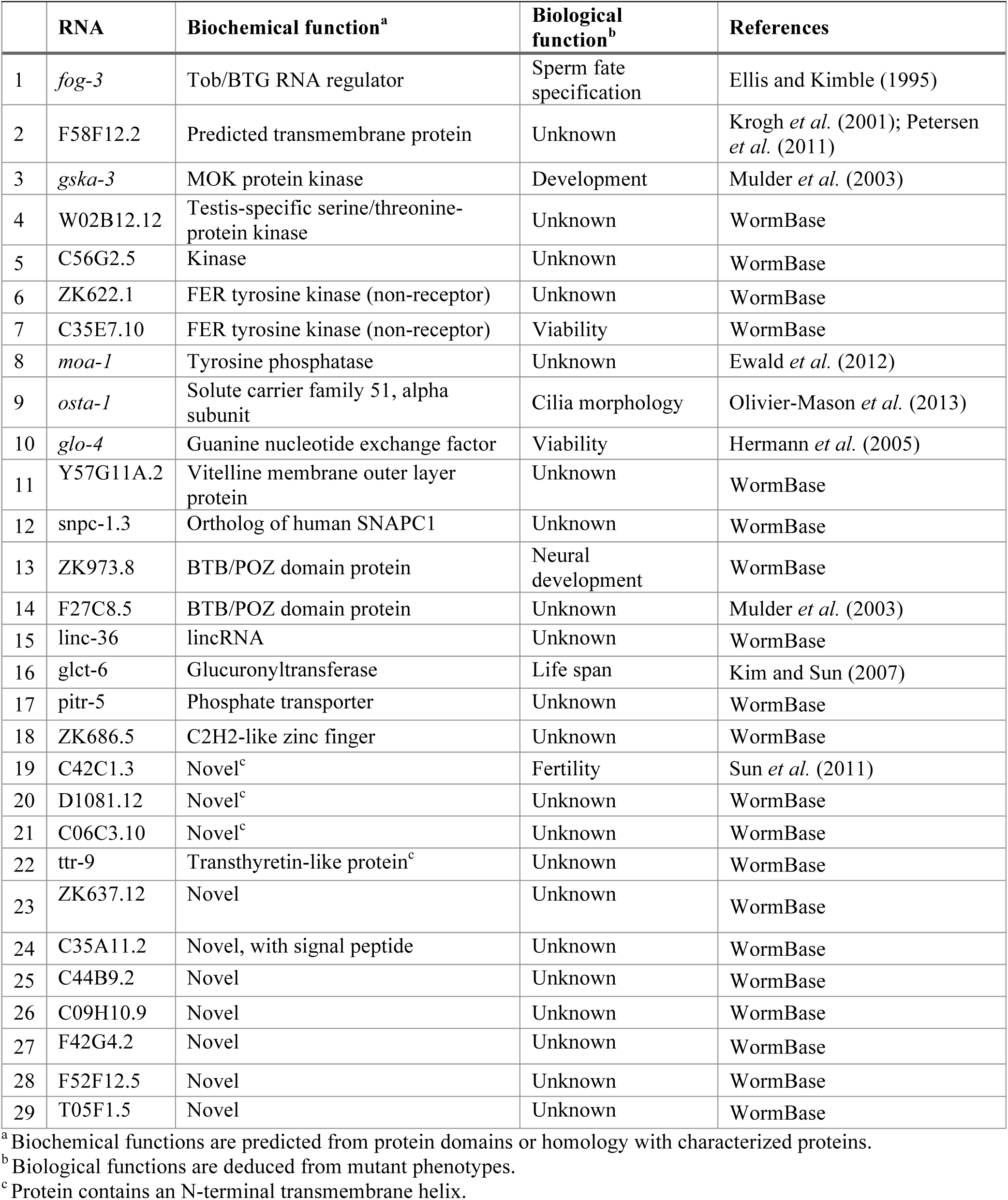
Major spermatogenesis-specific FBF targets, from Block I

### Block II RNAs (File S6)

Block II contains 510 RNAs, most of which were found in FBF iCLIP RNAs common to spermatogenic and oogenic germlines (Figure 3A). Importantly, among target RNAs, Block II RNAs account for half of all FBF iCLIP reads and hence for half of all FBF interactions (Figure 4B). The Block II cluster includes 10 validated FBF target RNAs (*fog-1, syp-2, fem-3, gld-3, htp-2, mpk-1, him-3, lip-1, fbf-1*, and *fbf-2*). Stem cell maintenance is the major FBF function and this function is not gamete-specific (Crittenden *et al.* 2002). Consistent with the idea that Block II RNAs might be central to stem cell maintenance, they include key self-renewal regulators *fbf-1* and *fbf-2* (Crittenden *et al.* 2002), and are most enriched for the biological process GO terms of cell cycle (p-value 10^-20^), cell division (10^-18^), and mitotic nuclear divison (10^-17^), embryo development ending in birth or egg hatching (10^-53^) and reproduction (10^-27^, all GO terms in File S3). Cell cycle regulation is central to stem cell maintenance (e.g. Orford and Scadden 2008), and we suggest that Block II is enriched in RNAs belonging to the FBF subnetwork responsible for stem cell maintenance.

### Block III RNAs (Table 2, File S6)

Block III contains 21 RNAs that are common to FBF iCLIP from spermatogenic and oogenic germlines, similar to Block II RNAs (Figure 3A). Block III RNAs stand out from Block II RNAs by their much higher frequency of FBF binding (Figure 4A). Because the frequency of RNA-protein complexes is a function of both affinity and abundance, we expected Block III RNAs to be abundant and to possess canonical FBF binding sites. Indeed, all Block III RNAs were abundant (Figure S4) and all had at least one canonical FBE under its highest peak: 15/21 had two or more canonical FBEs under that peak and 17/21 had an FBE with −1 or −2 C (known to enhance affinity) under that peak. Therefore, Block III RNAs emerge as exceptionally frequent FBF interactors due to both RNA abundance and high affinity FBF binding.

**Table 2.**
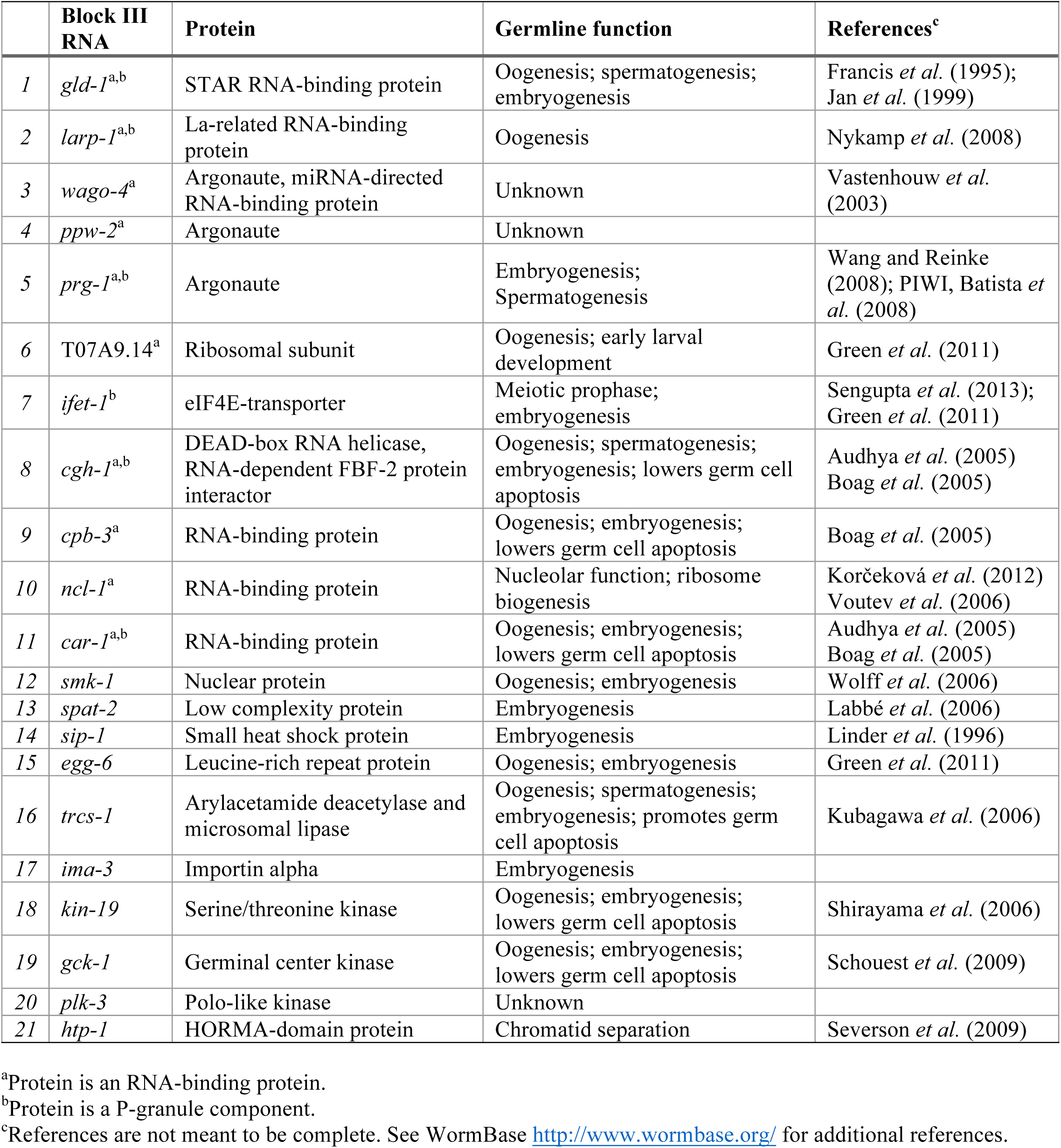
Block III RNAs, their protein products and germline roles

Block III RNAs also stand out by molecular function. Most striking is that 10/21 encode RNA regulatory proteins and 6/21 localize to P-granules (Table 2). GO terms (File S3) included P granule (10^-5^) and negative regulation of translation (<0.01). Block III includes two previously validated targets, *htp-1* and *gld-1*, the latter of which encodes a STAR RNA binding protein that localizes to P-granules, functions as a translational repressor and promotes differentiation (Francis *et al.* 1995; Jan *et al.* 1999; Jones *et al.* 1996). In addition, 3/22 were protein kinases (Table 2). The association of these molecular functions with Block III mRNAs, and hence with exceptionally frequent FBF targets, emphasizes the role of FBF as a regulator of other regulators and in particular, a high-level regulator of other post-transcriptional regulators.

Based on various functional studies, 16/21 Block III RNAs are required for gametogenesis or embryogenesis (Table 2). Roles in oogenesis and embryogenesis are best documented, perhaps because they have been analyzed more intensively than spermatogenesis. GO terms for oogenesis and embryo development ending in birth or egg hatching were both significant (p-values 10^-6^ and <0.01, respectively) Remarkably, 6/21 Block III mRNAs affect germ cell apoptosis (Table 2), a homeostatic mechanism common to worms and mammals, and the GO term apoptotic process was enriched (p-value <0.01). This finding underscores an earlier finding that FBF regulates MAPK-driven apoptosis in the germ line (Lee *et al.* 2007), a function conserved with murine PUM1 (Chen *et al.* 2012). Thus, many Block III RNAs are key regulators of gametogenesis, strengthening the notion that FBF maintains stem cells by repressing differentiation-promoting mRNAs. In addition, three Block III mRNAs are likely to regulate niche signaling in addition to promoting differentiation. The stem cell niche in this system relies on Notch signaling to maintain stem cells (Kimble and Crittenden 2007). Two Block III mRNAs encode physically interacting proteins, CAR-1 and CGH-1, that repress Notch signaling (Noble *et al.* 2008). A third Block III mRNA encodes CPB-3, a predicted binding partner of CGH-1 (Audhya *et al.* 2005; Boag *et al.* 2005). An attractive idea is that FBF represses expression of CAR-1 and CGH-1 in germline stem cells to enhance niche signaling.

We suggest that Block III RNAs also belong to the FBF subnetwork responsible for stem cell maintenance. Among these mRNAs, FBF appears to promote stem cell self-renewal in part by enhancing niche signaling and in part by repressing differentiation.

### Block IV RNAs (File S6)

Block IV contains 24 RNAs and represents RNAs enriched in oogenic germlines over spermatogenic germlines (Figure 2A). Of the 20/24 Block IV RNAs catagorized by Noble *et al.* (2016), all were in the oogenic program and 18/20 were only in the oogenic program. Interestingly, Block IV includes the snoRNA *ZK858.10*, which, being a snoRNA, is not found in Noble *et al.*, but which might still be an authentic part of the oogenic program. Consistent with oogenesis-related functions, Block IV also includes RNAs for the yolk receptor RME-2 and the ABC transporter MRP-4, the latter of which is expressed in oocytes to attract sperm (Kubagawa *et al.* 2006). However, Block IV, like Block I, includes many uncharacterized genes and no GO terms were enriched. Block IV likely represents an oogenesis-specific subnetwork.

### Conservation of Block I-IV PUF targets

We compared the RNAs in Blocks I-IV with the PUM2 iCLIP dataset from human embryonic kidney cells, as done for oogenic and spermatogenic FBF datasets described above (Figure 2F). Most striking was the 60% overlap of Block III RNAs with PUM2 targets. Block I had the lowest overlap among the three blocks with only ∼10%. Of the 21 high-frequency gender neutral Block III targets, 15/21 had human orthologs and 9 were also targets of human PUM2: *ncl-1*/TRIM2, *ima-3*/KPNA1, KPNA4, and KPNA5 (three orthologous PUM targets), *larp-1*/LARP1, *cgh-1*/DDX6, *gck-1*/STK24, *ifet-1*/EIF4ENIF1, *car-*1/LSM14B, *cpb-3*/CPEB2 and CPEB4, and *kin-19*/CSNK1E, CSNK1D and CSNK1A1. 7/9 of these are either RNA-binding proteins (*larp-1, cgh-1, car-3, ncl-1* and *cpb-3*) or regulate RNA processes (*ima-3, ifet-1*), consistent with a role for PUF proteins as regulators of other RNA regulators.

### Conclusions

Our analyses delineate clusters of FBF-bound RNAs that likely represent FBF subnetworks for spermatogenic (Block I), oogenic (Block IV), and stem cell (Blocks II and III) functions. Clearly this is only a first step in understanding the diverse roles of FBF regulation. Because stem cell regulation is a conserved function of metazoan PUFs and many RNAs in the FBF stem cell subnetwork are also targets of human PUM2, a next focus should be to learn whether phylogenetically conserved targets are subject to PUF regulation across phyla, and if so, how and where they are regulated.

## ACKNOWLEDGEMENTS

We thank Laura Vanderploeg for assistance with figure preparation, as well as Marie Adams and staff at the UW Biotechnology Center (UWBTC) for Illumina sequencing. We thank Anne Helsley-Marchbanks for assistance with preparation of the manuscript. This work was supported by NIH grants 5T32GM00869217 (AP), 5T32GM08349 (DFP), GM050942 (MW) and GM069454 (JK). JK is an investigator of the Howard Hughes Medical Institute.

## Supplemental figure captions

**Figure S1.**
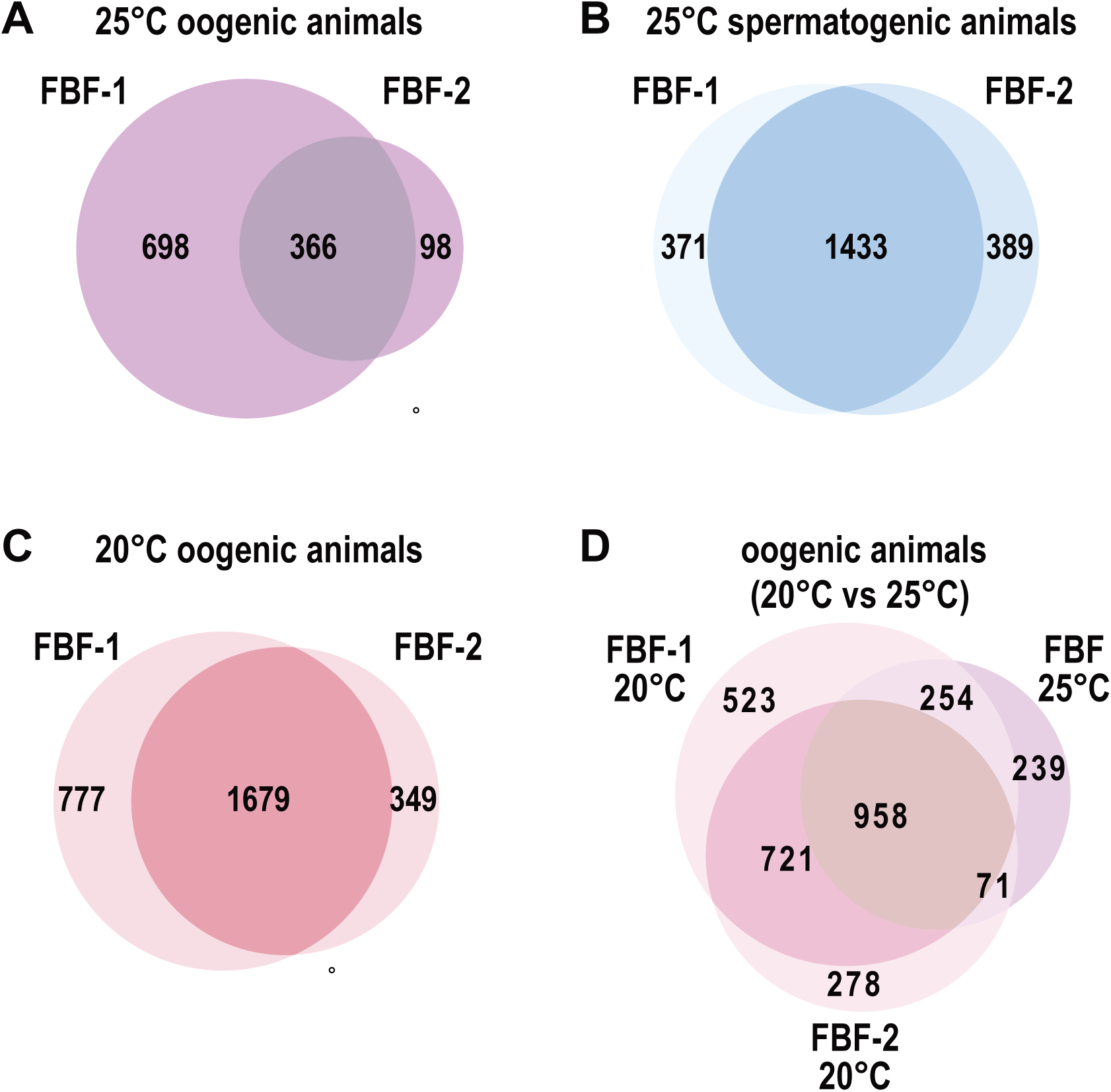
Venn diagrams of FBF-1 and FBF-2 RNA targets determined by iCLIP under different conditions. The two FBF proteins bind many RNAs in common, but also have individual targets. Numbers refer to the number of RNAs identified as FBF targets by iCLIP. (A) FBF iCLIP in oogenic worms raised at 25°. (B) FBF iCLIP in spermatogenic worms raised at 25°. (C) FBF iCLIP in oogenic worms raised at 20°. (D) FBF-1 and FBF-2 combined (“FBF 25°”) targets from oogenic worms raised at 25° compared to individually determined FBF-1 and FBF-2 targets from oogenic worms raised at 20°. The combined FBF-1 and FBF-2 (25°) target list overlaps equally well with the individual FBF-1 and FBF-2 (20°) target lists. This indicates that the combined FBF-1 and FBF-2 target set represents an average of FBF-1 and FBF-2.

**Figure S2.**
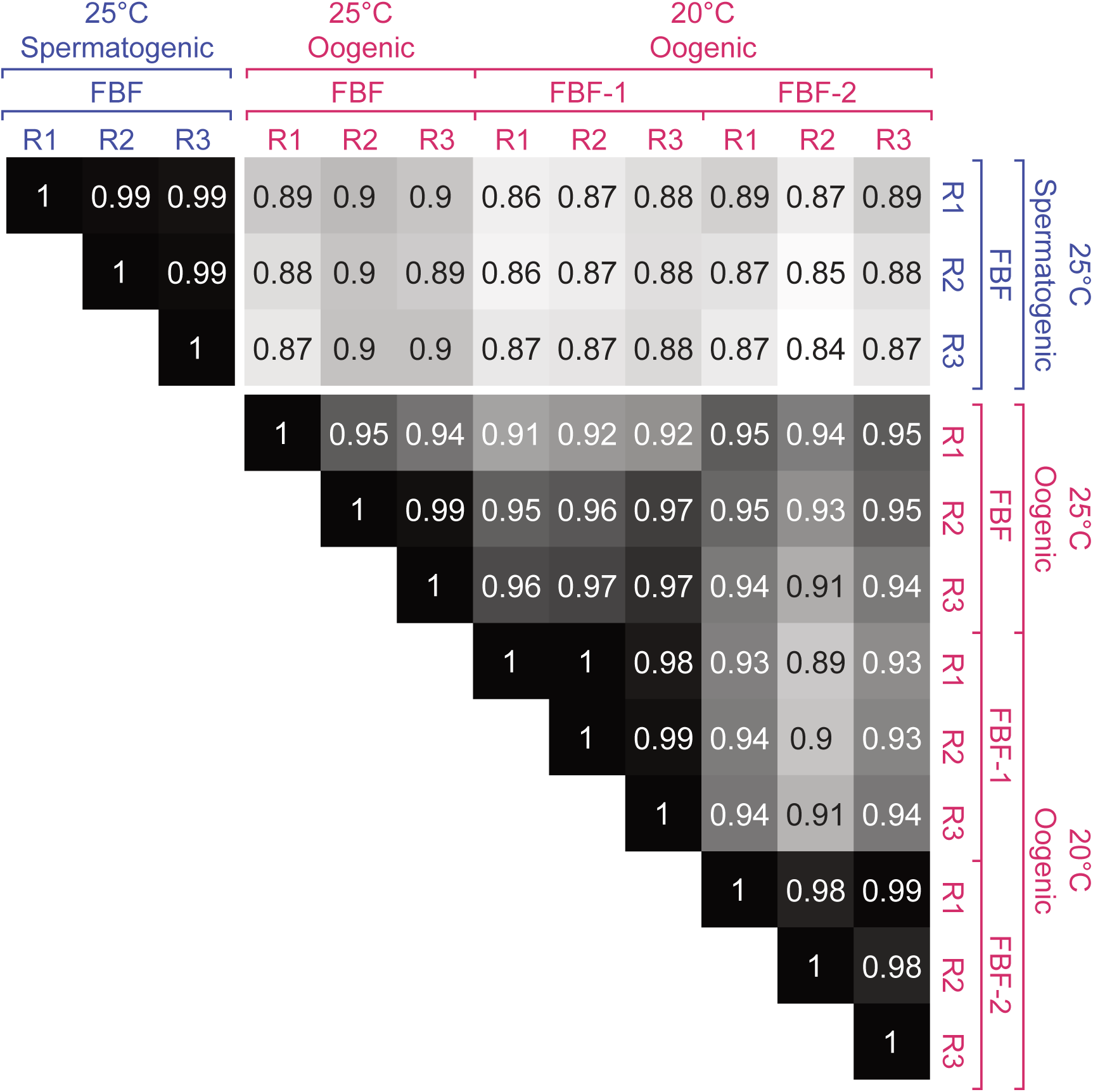
Spearman correlations between all iCLIP replicates (R) reinforce the conclusion that FBF has distinct binding landscapes in spermatogenic and oogenic germlines. Numbers represent rho values for the Spearman correlation between three replicates (R1 – R3) of iCLIP reads mapping to every target RNA. Target RNAs are defined as the 3,478 RNAs possessing a significant peak in any of the FBF iCLIP replicates. We used all RNAs identified as targets in any experiment to include all possibly relevant RNAs. Reads per gene were normalized to dataset size. 25° replicates represent combined FBF-1 and FBF-2 replicates (see Materials and Methods). The 25° oogenic FBF replicate (R1) was of lower complexity than the others, which likely explains its lower correlations overall.

**Figure S3.**
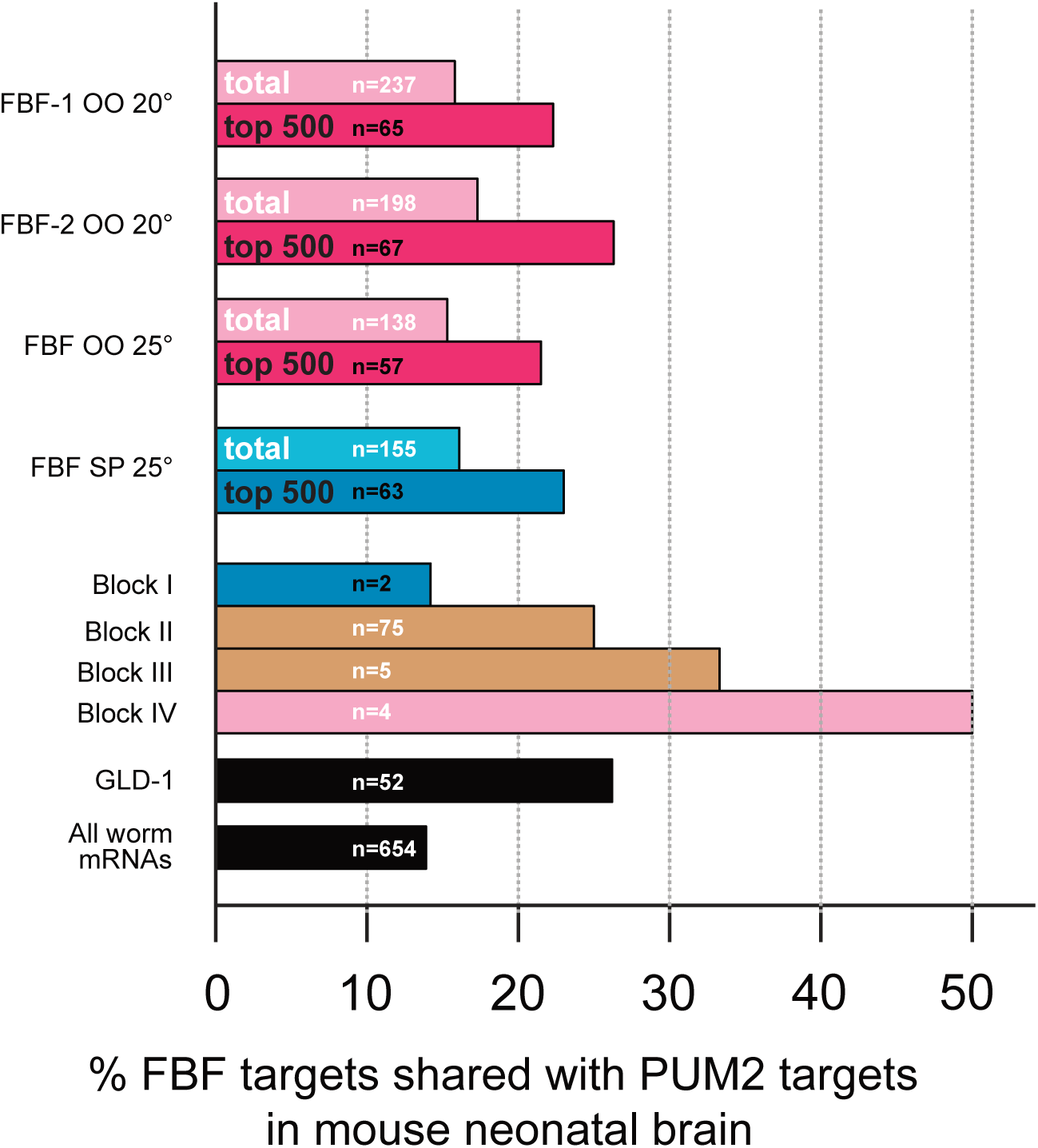
Overlaps of FBF iCLIP targets with human PUM1 and PUM2 iCLIP targets from murine neonatal brain (Zhang *et al.* 2017). As in Figure 2F, the number of ortholog groups comprising the overlap is indicated as “n=”. Overlap between GLD-1 targets of (Jungkamp *et al.* 2011) and PUM2 targets (Zhang *et al.* 2017) are included for comparison.

**Figure S4.**
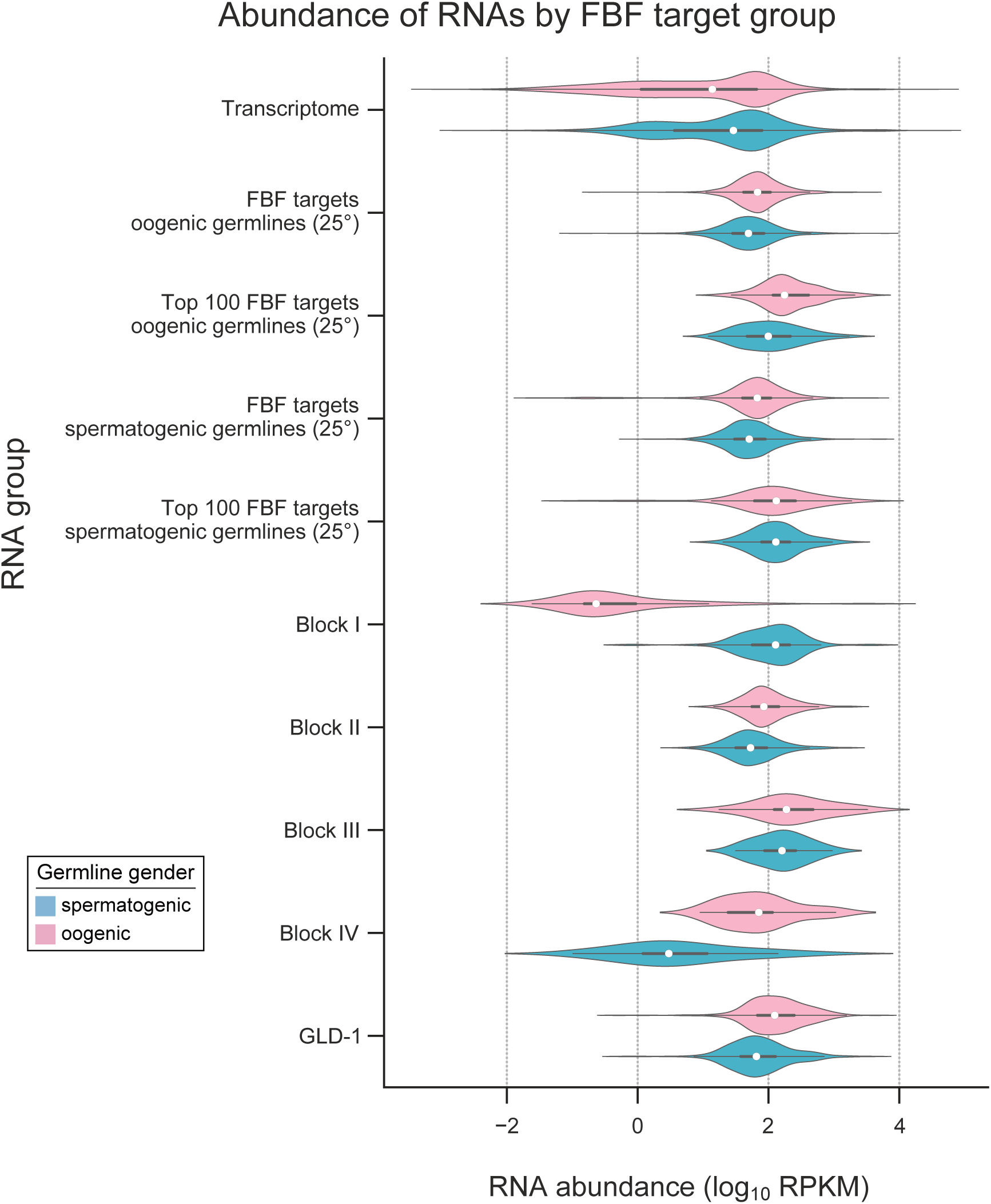
Top FBF targets are relatively abundant. The x-axis represents RNA abundance in log_10_ RPKM values for oogenic adult hermaphrodite gonads (Ortiz *et al.* 2014). On the y-axis, from top to bottom are: all RNAs present in the oogenic program (Noble *et al.* 2016), all targets of FBF in oogenic germlines (25°), the top 100 (by peak height) targets of oogenic (25°) FBF, and Blocks I-IV from Fig 3. The violin plot represents a Gaussian kernel density estimate fit to the data. An interior boxplot is also plotted: the white dot represents the median of the distribution, and the box indicates quartiles.

## Supplementary Files

**File S1.** Peaks called after FBF iCLIP from oogenic (oo) or spermatogenic (sp) animals at either 25° or 20°. Each tab label indicates germline gender, temperature and which FBF paralog was used for iCLIP. For the 25° datasets, FBF-1 and FBF-2 peaks are listed separately and shown as a combined list termed “FBF”, as described in Materials and Methods.

**File S2.** Metrics for FBF iCLIP peaks. Percentages of peaks with a canonical FBE are provided for both the total list and for the top 500 peaks, as defined in File S1 under the column labeled “Rank” (column “A”).

**File S3.** GO terms for FBF targets from all datasets, as well as Blocks II and III. Terms were identified using DAVID (Huang *et al.* 2009), and, except for Blocks I-IV, only the top 500 ranking RNAs in each dataset were included. The “Benjamini” column denotes the Benjamini-adjusted p value output by DAVID. There were no significant GO terms (p value < 0.01) for Block I or IV.

**File S4.** Genes significantly differing between spermatogenic and oogenic (both at 25°) iCLIP by DESeq2. Tab 1: Genes 2-fold enriched in spermatogenic iCLIP at P < 0.01. Tab 2: Genes 2-fold enriched in oogenic iCLIP at P < 0.01. Tab 3: all DESeq2 results.

**File S5.** FBF binding per gene for 2,114 FBF target RNAs. This dataset corresponds to Figure 3A.

**File S6.** Blocks, as defined in Figure 3.

**File S7.** FBF targets overlapping with the human PUF protein PUM2 identified by PAR-CLIP in HEK293 cells (Hafner *et al.* 2010). Tab names correspond to peak lists in File S1 or the blocks in File S6, except the “Common” tab corresponds to targets shared by FBF in both spermatogenic and oogenic germlines. Tabs labeled “top 500” only include the top 500 FBF targets in their respective list, ranked by frequency (read count). Worm locus IDs and gene names correspond to FBF targets. All human ortholog Ensembl IDs are given for each RNA, regardless of whether they are targeted by PUM2. The “PUM2 target HGNC symbol” column denotes orthologous PUM2 targets.

